# Steering Vector Fields for Property-Controlled Molecular Generation with Chemical Language Models

**DOI:** 10.1101/2025.09.24.678080

**Authors:** Aleksandar Dimitrievikj, Jude Wells

## Abstract

Chemical language models have recently become a powerful tool for the de novo generation of drug-like molecules represented as SMILES strings. A central challenge is steering generation toward compounds with favorable properties such as solubility and absorption. To this end, we investigate inference time control of generative chemical language models using activation steering. Using contrastive activation addition, we seek to improve three relevant properties: molecular size, aqueous solubility (log S), and lipophilicity (log P) without changing the model weights. We compare two interventions: a single global vector which is added to the activation in the last transformer layer, and a novel vector field where the addition vector is computed as a function of the current hidden state. Across multiple protein targets and two pre-trained models, the global steering vector yields desired results in just over half of our experiments, while the vector field achieves larger shifts at the expense of a decrease in the validity rate.

## 1 Introduction

Controlling molecular properties during de novo generation is a core challenge in computational drug discovery. Existing approaches such as policy-gradient reinforcement learning, conditional generation via property tags, and latent-space optimization can be compute-intensive and brittle under distribution shift or noisy property oracles [4, 11, 6, 5]. We investigate a lightweight, inference-time alternative—contrastive activation addition (CAA)—based on *activation steering* [8] for chemical language models (CLMs). CAA pre-computes a steering vector in the transformer activation space from contrastive pairs of positive and negative samples. During generation, this vector is added to the model’s activation state, nudging the distribution of generated molecules toward the desirable properties represented by the positive samples.

In contrast to post-hoc optimization pipelines, where conditional models must be retrained on property labels [1, 7] or reinforcement learning updates are performed after each generation step [4, 11, 6], activation steering offers a single inference-time knob with modest computational overhead. The traditional steering vector approach computes a *global steering vector* from contrastive molecule pairs, then adds this vector into a transformer layer during decoding. We also introduce a *token-aligned steering vector field* that conditions the added vector on the evolving hidden state at each generation step (figure 1). These interventions require only the hidden-state activations of a frozen CLM and do not modify the model weights.

**Figure 1:**
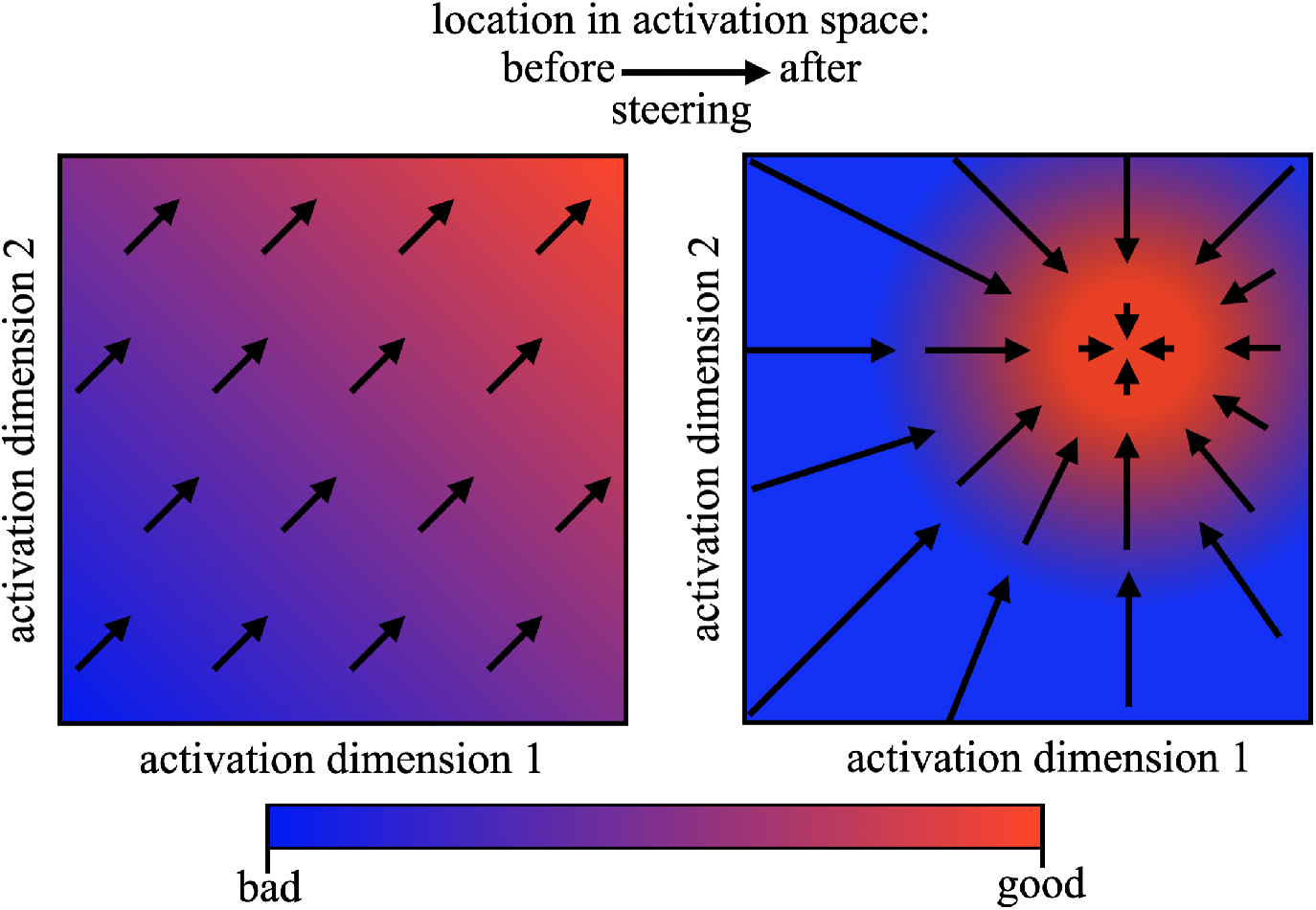
We depict a 2-D activation space where red denotes desirable regions and blue denotes undesirable ones. Left (global steering): a single constant vector is added everywhere, inducing a uniform vector field. The adjustment has the same direction and magnitude no matter where the unsteered activation starts. Right (vector-field steering): the steering vector depends on the current activation, so both direction and strength vary with location. This allows context-dependent adjustments that move states toward the desirable region.

We target three physico-chemical properties: molecular size (number of heavy atoms), aqueous solubility (logS), and lipophilicity (logP). These are inexpensive to predict using RDKit for atom counts, SolTranNet for solubility [3], and the Wildman–Crippen fragment model for logP [13].

Our study addresses two questions: (i) Can CLMs be steered toward desired property ranges via contrastively derived activation vectors? (ii) What are the trade-offs between property shift, validity, and uniqueness when applying a global vector versus a token-embedding-aligned vector?

### Contributions

(1) We adapt contrastive activation addition (CAA) to chemical language models and demonstrate decode-time control of size, log *S*, and log *P*. (2) We introduce a novel approach to activation addition steering: the steering vector field. (3) We compare the original global steering vector approach with our vector field and characterize the power–stability trade-off.

## 2 Methods

### Overview

We study test-time control for chemical language models (CLMs) via *contrastive activation addition* (CAA). CAA builds a “steering” direction in transformer-layer activation space from contrastive example pairs and injects that direction during decoding, leaving all model weights frozen. To compute the global steering vector, we proceed in three steps. First, we sample molecules from the model and construct positive–negative pairs based on the property of interest. Second, we run each sample through the model and record the final-layer activations at a specified token position. Finally, we compute the steering vector as the average of all difference vectors pointing from the negative to the corresponding positive sample across pairs. The global vector approach assumes that the desired shift in activation space has a constant direction and magnitude, regardless of the current location. This assumption is limiting for properties such as logP, where successful steering requires values to fall within a desirable range rather than being maximized or minimized. To address this, we introduce a variant of CAA that we call a steering vector field. Here, the steering vector is defined as a function of the current activation state. Instead of using a global mean of all difference vectors, the field computes a weighted mean in which each pair contributes proportionally to its proximity: vectors from negative samples closer to the current activation state receive higher weight. We evaluate both approaches on two CLMs, DrugGen [10] and Plixer [12].

### 2.1 Generative model

#### DrugGen

DrugGen [10] is a GPT-style CLM fine-tuned on ∼800k (target, SMILES) pairs. Using it, we sample molecules using the prompt `<|startoftext|><P>{protein_sequence}<L>`

#### Plixer

Plixer [12] is a two-stage pocket-conditioned generator: (i) *Poc2Mol*, a 3D U-Net that inpaints ligand voxels inside a protein-pocket voxel grid from a PDB structure; (ii) *Vox2Smiles*, a ViT encoder with a GPT-style decoder that converts ligand voxels to SMILES. We steer only the decoder of Vox2Smiles at layer *ℓ* = 11; weights stay frozen.

#### 2.2 Protein targets and dataset curation

##### DrugGen

We uniformly sampled 21 UniProt targets from the training set (list in Appendix). For each UniProt ID we retrieve the canonical sequence and prompt DrugGen to generate an initial pool of *N* = 250 distinct SMILES. Invalid strings are rejected with RDKit; the remainder define the per-target baseline.

##### Plixer

We use 10 protein pockets with resolved co-crystal ligands. For each pocket we compute the pocket center from the ligand, remove the ligand from the PDB, voxelize the pocket with the repository defaults, and run the combined protein-to-SMILES pipeline to produce an initial pool of *N* = 250 distinct SMILES. Invalid strings are rejected with RDKit; the remaining unique molecules define the per-target baseline.

### 2.3 Constructing contrastive pairs

Given valid molecules 𝒮 = {*s*_1_, …, *s*_*m*_} for protein *P*, we form property-specific pairs:

1. **Size** (heavy-atom count *n*_at_): create ordered pairs (*s*_*i*_, *s*_*j*_) with 0.6 *n*_at_(*s*_*j*_) ≤ *n*_at_(*s*_*i*_) ≤ 0.8 *n*_at_(*s*_*j*_), marking the smaller molecule as positive.
2. **Aqueous solubility** (log *S*): predict with SolTranNet [3]; in a pair, the more soluble molecule is positive.
3. **Lipophilicity** (log *P*): estimate via Wildman–Crippen fragments; positive if 1 ≤ log *P* ≤ 3, negative otherwise, following the recommended drug-like window [2].

Each element is prefixed with the protein prompt so the model sees complete sequences during activation capture and 50 molecules are generated as a pool of molecules for constructing the pairs.

### 2.4 Contrastive Activation Addition (global vector)

Let **h**^(*ℓ*)^(*x, t*) *∈* ℝ^*d*^ be the hidden state at layer *ℓ*, token *t* for string *x*. With positive set 𝒫 and negative set *N* (from §2.3), the *global steering vector* at layer *ℓ* is

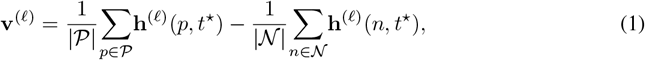

where *t*^*⋆*^ is a tunable token index (we use the last token before `<eos>` unless stated otherwise; cf. [9]). At inference time we apply a linear intervention to all SMILES tokens,

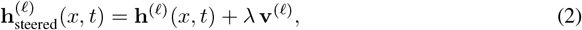

*λ* controls effect size.

### 2.5 Token-aligned steering vector field

A single **v**^(*ℓ*)^ applies the same offset everywhere. To capture location-dependent variation in steering direction, we build a state-dependent *vector field*. For each pair (*n*_*j*_, *p*_*j*_) we store

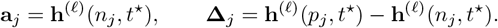

At decode step *t*, let *T*_seq_(*x*) be the current sequence length and

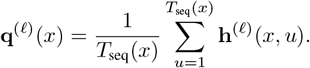

We compute weights with a temperature-scaled softmax over *ℓ*_2_ distances:

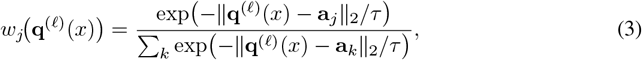

and define the local direction

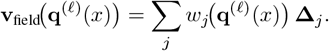

We intervene as 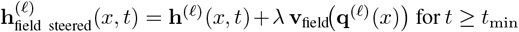. Smaller *τ* concentrates on nearest anchors (stronger but less stable); larger *τ* smooths updates. This induces a bias–variance trade-off: global vectors are stable but under-correct local structure; vector fields give larger shifts but can reduce validity if too sharp.

### 2.6 Experimental protocol

For each protein: (i) build a steering object (global vector or vector field) from |𝒫| unique molecules and the chosen (ℓ, *t*^*⋆*^); (ii) generate *G* = 250 molecules under two conditions: no steering (*λ* = 0) and positive (+*λ*), (iii) log summary stats and save SMILES.

### 2.7 Evaluation metrics

We compute *molecule-level* validity, uniqueness, and whether it is identical to a generated molecule used for computing the steering vector (pair-pool-leaked), then test *target-level* property shifts (mean size, mean log *S*, in-range fraction for log *P*). Results are reported as #successes/#eligible targets per property and intervention.

## 3 Results

### Results at a glance

For around half of our experiments across protein targets, properties and models we see a clear and consistent shift in the desired direction. In most of these successful cases, we even see that the positive tail of the distribution for size and log *S* is shifted along with the mean, showing that steering can ‘unlock’ parts of molecular space that could not have been reached merely from oversampling the non-steered model (figures 3 and 4). However, we have to acknowledge that for many targets, steering did not have a significant effect; in some cases, the effect was opposite to the intended direction (figure 2). Overall, this suggests that steering is model and protein-target dependent. It is worth trying if the results can be quickly and cheaply evaluated (as is the case with our computationally assessable properties), but it is not reliable enough to assume steering will have the desired effect for an arbitrarily chosen target.

**Figure 2:**
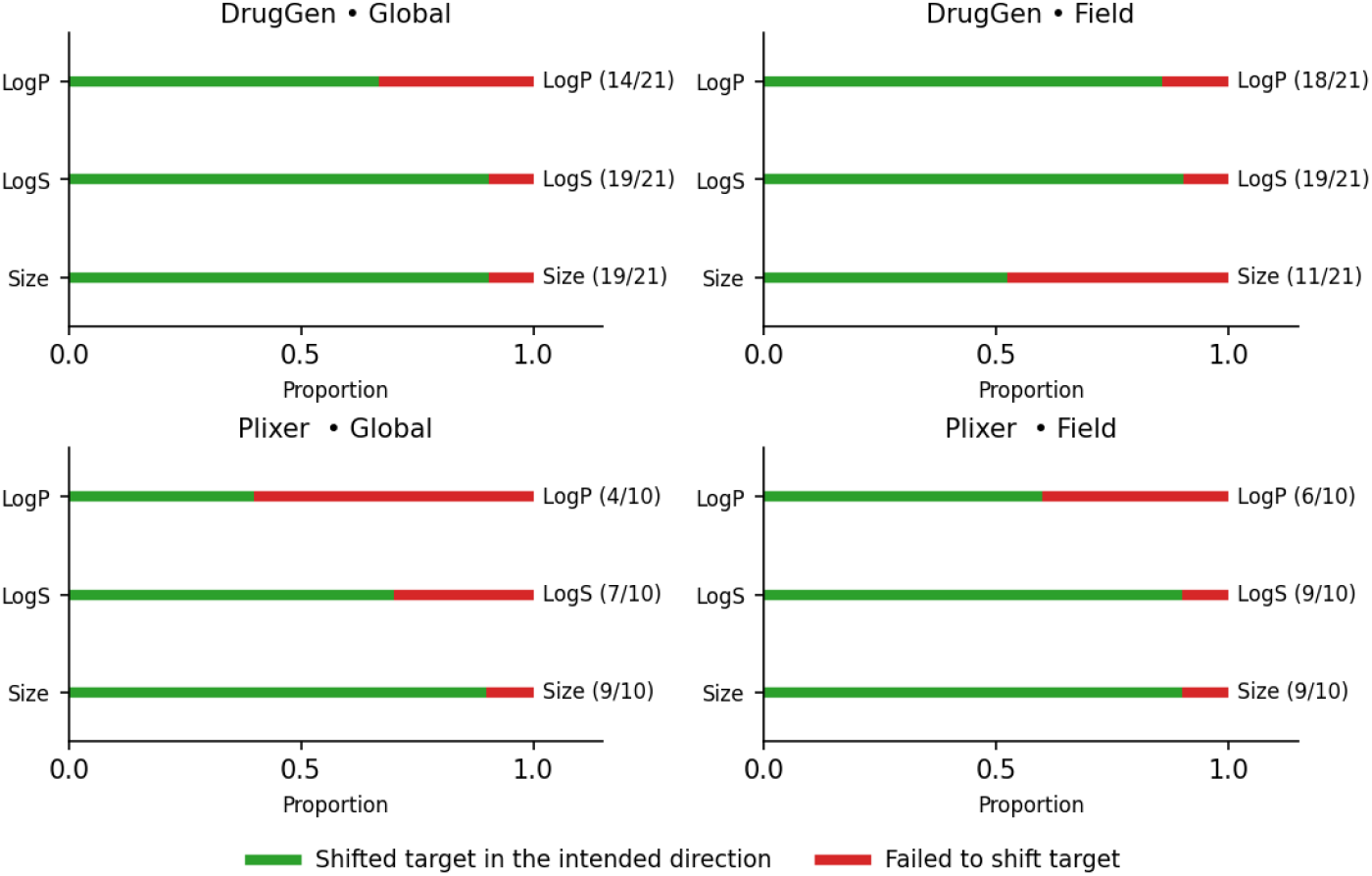
Unfiltered directional shift rates. For each model and intervention, bars show the proportion of targets where the mean property moved in the intended direction (no significance test and no quality filter). These rates are higher than the filtered success counts in Table 1 because they ignore validity/uniqueness thresholds and p-value criteria.

**Figure 3:**
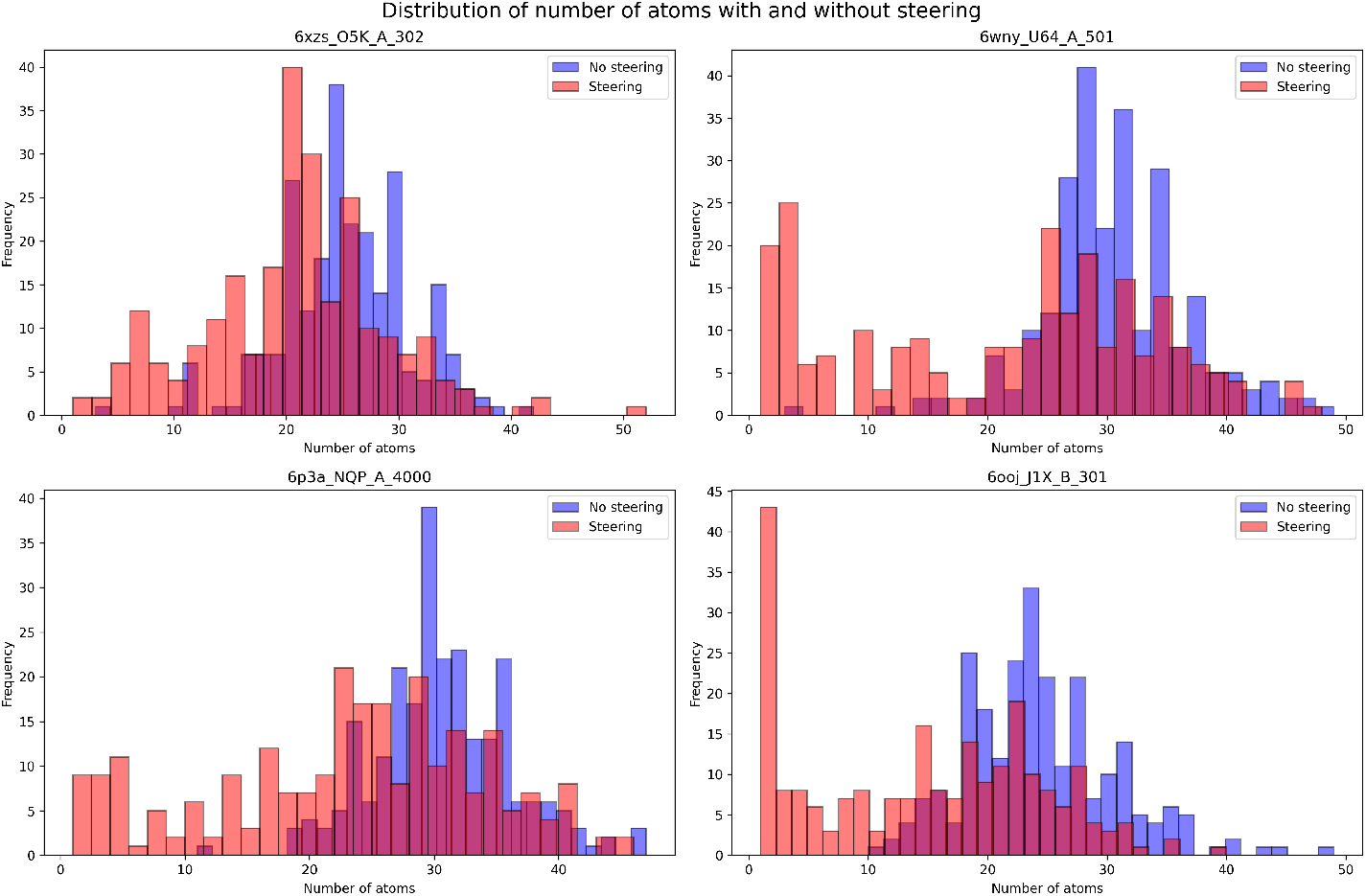
Overlaid histograms of heavy-atom counts for four representative targets using the field intervention on Plixer (baseline blue vs. steered red). Steering shifts the mass of the distribution toward smaller molecules while preserving the overall shape for most targets. These examples are drawn from targets that pass the baseline quality filter.

**Figure 4:**
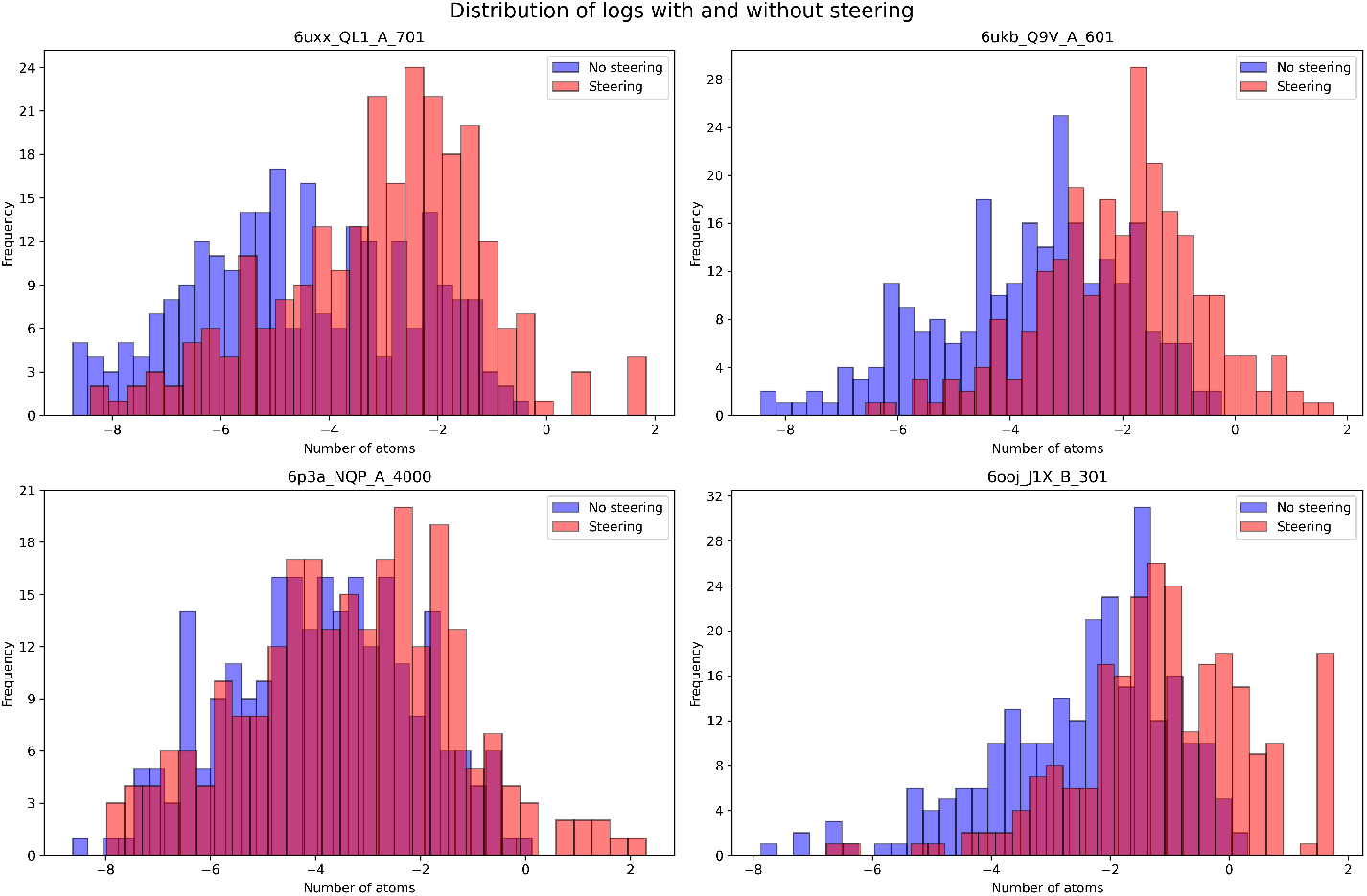
Overlaid histograms of log *S* predictions for four representative targets using the field intervention on Plixer (baseline blue vs. steered red). Steering shifts the distribution toward higher predicted solubility on most targets. Examples shown pass the baseline quality filter.

### 3.1 Success criteria and filtering

A target counts as *successfully steered* for a property only if **both** hold:

i. the intervention effect is in the correct direction and statistically significant on the *filtered* set; and
ii. at least 10% of generated molecules are *valid, non-duplicate*, and *not pair-pool-leaked*.

We pre-exclude targets whose *baseline* pool fails (ii). For size and log *S*, significance is a two-sided Welch *t*-test on the filtered molecules (*p<*0.05). For log *P*, the outcome is the *proportion in range* [1, 3]; we test whether this proportion is higher under the intervention using a one-sided two-proportion test (Fisher’s exact test when any expected count is *<* 5), with success at *p<*0.05.

#### Multiple testing

P-values are computed per target and are not corrected across targets; the counts we report are descriptive robustness indicators rather than family-wise error claims.

### Main results table

Each cell reports `#targets passing / #eligible targets` for a model (rows), property (blocks), and intervention (**Global** vs. **Field**). Only targets that meet the baseline quality filter (ii) are counted as eligible.

### Directional tendency vs. filtered success

Figure 2 summarizes raw steerability, whether the mean shifted in the intended direction without applying the eligibility filter or significance tests. As expected, these directional rates exceed the filtered success counts in Table 1. The gap quantifies where steering nudges the distribution but either (i) effects are small and fail the statistical test, or (ii) validity/uniqueness losses reduce eligibility.

**Table 1:** Fraction of targets passing the success checks per property and intervention.

|  | size |  | log(S) |  | log(P) |  |
| --- | --- | --- | --- | --- | --- | --- |
|  | Global | Field | Global | Field | Global | Field |
| DrugGen | 15/21 | 7/21 | 14/21 | 17/21 | 9/21 | 13/21 |
| Plixer | 6/10 | 8/10 | 4/10 | 8/10 | 1/10 | 3/10 |

### Overall patterns

DrugGen: the **Global** vector is strongest for *size* (15/21) and competitive on log *S* (14/21), while the **Field** dominates on log *S* (17/21) and log *P* (13/21). Plixer: the **Field** outperforms **Global** on all three properties (size 8/10 vs. 6/10; log *S* 8/10 vs. 4/10; log *P* 3/10 vs. 1/10). These counts indicate a power–stability trade-off: the field achieves larger shifts more often; the global vector is more conservative.

### Model comparison

Plixer benefits more from the **Field** than DrugGen across all properties, consistent with a decoder that conditions on richer intermediate representations, where a single global direction under-corrects local structure. DrugGen shows a safer profile for size with the **Global** intervention.

### Size distributions

Figure 3 shows a reduced heavy atom count under steering both in the mean and the left tail.

### Solubility distributions

Figure 4 shows right-shifted log *S* histograms under steering. Several targets exhibit heavier right tails, suggesting the steered model is generating molecules that would be extremely unlikely under the non-steered model.

### Lipophilicity

For log *P* we summarize via in-range counts only (Table 1). Gains concentrate on a subset of targets and favor the **Field** intervention on both models.

### Quality effects

Validity and uniqueness remain close to baseline for **Global** on DrugGen; the **Field** can reduce validity on harder targets, consistent with stronger, state-dependent updates. These effects are reflected in eligibility and pass counts.

## 4 Discussion

Our results show a clear trend suggesting that both Global CAA and steering vector fields can be used to bias a range of molecular properties without having to fine-tune the generative model. However the effectiveness is dependent on the protein target and desired property, working very well for some proteins and failing on others. For some combinations of steered property and model our vector field approach outperformed the global vector but it was not consistently better. The flexibility of the field comes with risks: when the current hidden state lies outside the anchor cloud, interpolation extrapolates, pushing decoding off-manifold and reducing validity. Compute-wise, the field adds a lightweight per-step lookup and weighted sum over anchors; the overhead scales with anchor count. We did not extensively explore hyper-parameters such as intervention layer token index and *τ* (the softmax smoothing parameter for the field) however we did find it essential to adjust *λ* (multiplicative factor on the added vector) when switching between models. Our observations suggest that the magnitude of the steering vector is a very important factor: too low will limit the effect size, while too high can yield invalid generations.

## 5 Limitations

- **Proxy oracles**. Solubility and lipophilicity are estimated by predictors (SolTranNet, fragment log *P*); steering may exploit non-physically meaningful aspects of these oracles.
- **Vector field**. Performance can degrade when the current state lies outside the anchor coverage; the interpolated direction then extrapolates and may reduce validity.
- **No retraining baselines**. We do not compare against retrained conditional or RL baselines here; results show feasibility of test-time control, not superiority.
- **Hyperparameter sensitivity**. Meaningful shifts require tuning of layer *ℓ*, token index *t*^*⋆*^, steering scale *λ*, and (for the field) temperature *τ*. The best settings differ by model and property; naïve transfers can underperform or hurt validity.

## Acknowledgments and Disclosure of Funding

Jude Wells acknowledges the receipt of studentship awards from the Health Data Research UK-The Alan Turing Institute Wellcome PhD Programme in Health Data Science (Grant Ref: 218529/Z/19/Z).

## A Hyperparameters for steering

Unless otherwise noted, we use the following fixed settings:

- **Layer and token index**. Late transformer layer *ℓ* = 11; token index *t*^*⋆*^ is the last token before the stop token.
- **Intervention scope**. Additions are applied to all SMILES tokens
- **Steering scale**. *λ* = 1 for positive steering for DrugGen and *λ* = 3 for Plixer (global variant only)
- **Vector-field temperature (if used)**. *τ* = 0.1 for DrugGen and *τ* = 1 for Plixer.
- **Pair construction pool**. Per target, construct contrastive pairs from a pool of *M* = 50 generated molecules, avoiding duplicates when required.
- **Sampling**. For each target, generate *G* = 250 molecules per condition.
- **Start of SMILES**. *t*_min_ is the first SMILES token; all interventions apply only for *t ≥ t*_min_.

## B Protein targets

### DrugGen

The 21 UniProt ids used in the DrugGen experiments:

~~~
P07900, P00734, Q14524, P19823, P78334, P14416, P08913, P03372, P35348,
P08172, P09622, P36956, P20309, P04818, P25100, P18825, P23634, Q8N1C3,
P03372, P48169, P10275
~~~

### Plixer

The 10 pockets used in the Plixer experiments:

~~~
6wny_U64_A_501, 6zxo_D16_C_401, 6s4h_KUQ_A_901, 6xzs_O5K_A_302,
6ukb_Q9V_A_601, 7nqw_UNW_A_405, 6ooj_J1X_B_301, 6tx4_HRZ_A_204,
6p3a_NQP_A_4000, 6uxx_QL1_A_701
~~~

